# E6AP is important for HPV E6’s role in regulating epithelial homeostasis and its loss impairs keratinocyte commitment to differentiation

**DOI:** 10.1101/2022.06.23.497250

**Authors:** Wen Yin, Nagayasu Egawa, Ke Zheng, Heather Griffin, Ademola Aiyenuro, Jacob Bornstein, John Doorbar

## Abstract

Human papillomaviruses (HPV) typically cause chronic infections by modulating homeostasis of infected basal cell to ensure persistence. Using FUCCI and cell-cell competition assays, we established the role of two common viral targets of low-risk and high-risk E6 proteins, E6AP and NHERF1, on four key components of epithelial homeostasis. These includes cell density, proliferation, commitment to differentiation and basal layer delamination. Our RNA sequencing results validated E6’s effects on homeostasis and revealed similar transcriptional gene regulation of E6-expressing cells and E6AP^-/-^ cells. For example, yes-associated protein (YAP) target genes were up-regulated by either E6 expression or E6AP depletion. This is also supported by YAP expression pattern in both monolayer cell culture and HPV-infected clinical tissues. As the conserved binding partner of Alpha group HPV E6 proteins, the precise role of E6AP in modulating keratinocyte phenotype and associated signalling pathways have not been defined. We demonstrate that deletion of E6AP in keratinocytes delayed the onset of differentiation and the abundance of E6AP is reduced in HPV-infected tissue. This suggests that Alpha E6 regulates epithelium homeostasis by inhibiting E6AP’s activity, leading to alteration of multiple downstream pathways including YAP activation. Potential treatments can thus be developed to resolve the reservoir of HPV infection.

## Introduction

Human papillomaviruses (HPV) are non-enveloped double-stranded DNA viruses that infect multiple epithelial sites (McBride, 2017; John Doorbar, 2015). So far, more than 200 HPV types based on L1 viral gene sequence identity have been discovered, which are classified into five genera: Alpha-, Beta-, Gamma, Mu and Nu-papillomaviruses (Bernard et al., 2010). The Alpha genus comprised of viruses that can either infect cutaneous or mucosal epithelial sites, those mucosal HPVs can be further classified into high-risk and low-risk HPVs based on the cancer risk associated with their infections (Doorbar et al., 2012; John Doorbar, 2015). HPV16 and HPV11 are representatives for the Alpha high-risk group and the low-risk group. HPVs generally cause self-limiting epithelial lesions that are usually resolved by the host immune system over time. However, the high-risk mucosal HPV infections can sometimes persist and lead to carcinomas (McBride, 2022). Although low-risk HPVs rarely cause malignancies, the size and location of the benign papillomas can render these lesions medically serious (Egawa & Doorbar, 2017). For example, recurrent respiratory papillomatosis (RRP) caused by HPV11 in children has no effective treatment and can only be controlled by repeat surgery. Condyloma acuminatum caused by HPV6 and HPV11 is one of the most widespread sexually transmitted disease (Goon et al., 2008; Ryan Ivancic et al., 2018).

The epidermis is a stratified epithelium composed of morphologically distinct cellular layers that reflect the terminal differentiation process of keratinocytes (Watt, 1989). Basal keratinocytes proliferate and expand to reach a specific density which triggers contact inhibition signals. This causes the cells to exit the cell cycle and commit to differentiation. Subsequently, keratinocytes leave the basal layer (delamination) and undergo an upward-directed transit into the more superficial spinous, granular, and cornified layers (Rice & Rompolas, 2020). Thus, epithelial homeostasis is maintained by the careful regulation of proliferation, basal cell density, delamination and differentiation. HPV viral proteins impart advantages to infected basal cells through modulating these key procedures, leading to lesion expansion and maintenance (Doorbar et al., 2021). Previous literature has indicated that E6 protein as a homeostasis regulator during productive infection, and in both the high and low-risk Alpha types, can regulate p53 and thus indirectly Notch-mediated epithelial differentiation (Khelil et al., 2021; Kranjec et al., 2017; Murakami et al., 2019; Yugawa et al., 2007). Moreover, accumulating evidence suggests that E6 modulates epithelial homeostasis through interacting with key molecules involved in WNT, NOTCH and HIPPO signalling (Gupta et al., 2018; Lichtig et al., 2010; Olmedo-Nieva et al., 2020; Sominsky et al., 2014). There is increasing evidence that the HIPPO pathway effector yes-associated protein (YAP) plays pivotal role in controlling epidermal homeostasis and high-risk E6 has been shown to control YAP nucleus-cytoplasm shift to activate YAP transcriptional activity (He et al., 2015; Webb Strickland et al., 2018). The HIPPO pathway senses mechanical cues such as cell density and contact signals from the basement membrane. Once activated, YAP is phosphorylated by LATS1/2 kinases and sequestered in the cytoplasm. When the hippo kinases are inactive, YAP is translocated into the nucleus and activate the downstream genes to drive keratinocyte proliferation (Corley et al., 2018; Webb Strickland et al., 2018). It was also shown that the inhibition of YAP activity triggers keratinocyte differentiation (Totaro et al., 2017; Totaro et al., 2018).

It has been reported previously that the Alpha genus E6 proteins are the only group that preferentially interacts with E6AP rather than MAML1, amongst which some E6 proteins further acquired the ability to induce E6AP degradation (Brimer et al., 2017). E6 typically associates with E6AP to form a complex that recruits secondary substrates such as p53 or NHERF1 for proteasomal degradation (Brimer et al., 2017; Drews et al., 2019; Zimmermann et al., 1999). Indeed, we believe that these cellular targets of E6 are important regulatory factors involved in epithelial homeostasis control. As the conserved binding partner of Alpha group E6, E6AP is likely to play a central role and the consequences of E6AP association vary among E6 proteins. For instance, E6AP binding does not necessarily trigger its ubiquitination activity (Brimer et al., 2017). Although the downstream consequences of E6AP degradation by E6 are poorly understood, it was reported that E6AP regulates WNT signalling, which is intensified by E6 in primary keratinocytes (Sominsky et al., 2014). Also, E6AP was found to promote cell growth in multiple cell types (Amit Mishra et al., 2009; Ramamoorthy et al., 2012; Srinivasan & Nawaz, 2011), indicating a potential role for E6AP to regulate skin homeostasis. NHERF1 is one of the first cellular targets discovered to be degraded by both high- and low-risk E6-E6AP complex, which has been shown to interact with beta-catenin and YAP directly (Accardi et al., 2011; Georgescu et al., 2016). Also, it was demonstrated that NHERF1 degradation by 16E6 led to activation of the WNT pathway (Drews et al., 2019). Thus, E6-directed NHERF1 degradation may also contribute to homeostasis regulation in epithelium.

Low-risk HPVs successfully survive and cause chronic lesions in the epithelium (Del Pino et al., 2012). Thus, low-risk HPVs must clearly possess the basic set of homeostatic functions to support their persistence in the basal epithelium (Doorbar et al., 2021). It is anticipated that these functions are conserved across the Alpha papillomavirus genus and important for maintaining virus lifecycle. In this study, we have dissected E6 functions in regulating homeostasis in normal spontaneously immortalised keratinocytes (NIKS) (Allen-Hoffmann et al., 2000) and found that E6AP and NHERF1 play important roles in these processes. Moreover, Clinical observations support the idea that key cellular targets of E6, including E6AP, NHERF1 and YAP, are regulated by the virus in the basal layer during HPV lifecycle. Intriguingly, we discovered, to our surprise, that E6AP has novel functions as a homeostasis modulator by using E6AP^-/-^ NIKS keratinocyte cell lines. RNA sequencing results revealed that the absence of E6AP suppressed the normal differentiation-related gene expression pattern in keratinocytes. Additionally, E6 expression or E6AP depletion activated a similar subset of YAP target genes. Therefore, our results suggest that E6 regulates key homeostatic processes in epithelium basal layer through inhibiting E6AP function. Disruption of the homeostatic pathway may have detrimental effects on viral persistence and offers attractive target for therapeutic approaches.

## Results

### 11E6 and 16E6 proteins modulate the balance between cell proliferation and differentiation

Recent work has demonstrated the role of E6 proteins in regulating multiple aspects of homeostasis and signaling pathways in keratinocytes. This includes E6-enhanced keratinocyte proliferation, E6-mediated inhibition in keratinocyte differentiation as well as effect on cell-cell contact (Herfs et al., 2017; Kranjec et al., 2017; Murakami et al., 2019; L Sherman & Schlegel, 1996; Levana Sherman et al., 2002). To maintain basal layer homeostasis, four processes need to be precisely controlled: cell cycle entry that suggests proliferative potential, cell density, timing of delamination and differentiation (Saunders-Wood et al., 2022). Firstly, to monitor the impact of E6 proteins upon cell cycle status, the fluorescent ubiquitination-based cell cycle indicator (FUCCI) system was used (Koh et al., 2017; Saitou & Imamura, 2016). FUCCI relies on the phase-dependent proteolysis of the oscillators Cdt1 and Geminin, and it is a powerful tool in visualizing cell cycle progression (figure 1A). By combining the FUCCI system with differentiation marker K10 and DAPI staining, this system allows the examination of three of the four components: cell cycle, differentiation and saturation density. Preliminary results using this system indicated that when FUCCI-expressing NIKS cells transduced with empty vector (EV) reached post-confluence, there were more Cdt1-mKO2 positive cells (red) and less Gemini-mAG positive cells (green) compared to E6-expressing NIKS cells at similar density (figure 1B). This is compatible with our understanding of E6-mediated cell proliferation upon contact inhibition. Also, NIKS-EV displayed higher K10 expression than NIKS-E6 cells, suggesting more cells committed to terminal differentiation in the absence of E6 expression. NIKS-E6 cells grow at different rates compared with NIKS-EV (Kranjec et al., 2017; Murakami et al., 2019), thus we decided to plate NIKS cells at pre-to post-confluence to assess the impact of E6 expression on cell phenotypes at different densities. Cells were left to adhere and form contacts for 72 hours after plating, then fixed and scanned by Confocal microscopy. High content imaging allows quantifications of the number of cells, K10-positive cells and Geminin-mAG positive cells for each field (figure 1C-D). In normal keratinocytes, as the cell density increases, contact inhibition is triggered and most cells enter G1 or G0 phase and begin to differentiate (Rice & Rompolas, 2020; Watt et al., 1988). Thus, the number of geminin-positive cells per field decreased as the number of cells increased. As shown in figure C, geminin-positive NIKS-EV cells declined from 32.6% to 7% and reached the steady state level. Cells started to express K10 after saturation density has reached approximately 10,000 cells per field (figure D). In comparison to NIKS-EV cells, the number of geminin-positive cells for 11E6-expressing NIKS was 37.7% at pre-confluence and dropped to 12% at post-confluence. For 16E6-expressing NIKS, geminin positive cells decreased from 46.4% to 14%. Thus, an increase of cycling cells was observed at both pre- and post-confluence for NIKS expressing either 11E6 or 16E6. Additionally, the expression of 11E6 or 16E6 increased the cell saturation density to approximately 14,000 cells or 17,000 cells/field respectively and delayed the threshold in which keratinocyte differentiation is triggered. Overall, our results suggest the roles of E6 in adjusting homeostasis steady state which alters the timing of cells transition from proliferation to differentiation.

**Figure 1.**
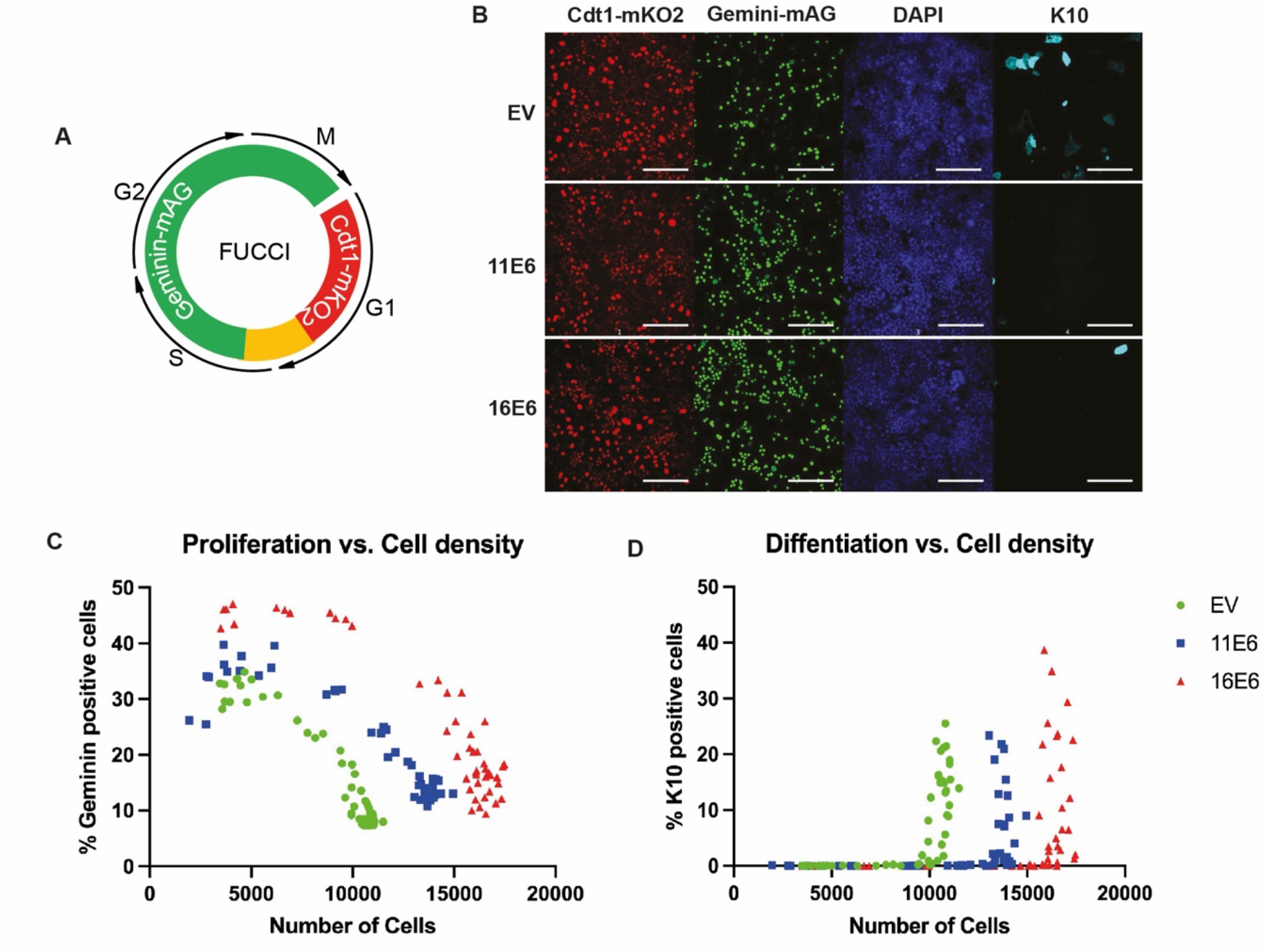
11E6 and 16E6 proteins regulate cell cycle progression, saturation density and differentiation of human keratinocytes. (A) A schematic diagram of FUCCI system is shown. (B) FUCCI-expressing NIKS cells were stably transduced with retroviral vectors harbouring empty vector (EV), 11E6 and 16E6 and cultured for 72 hours to reach post-confluence (10,000 cells/field). NIKS cells were fixed with 4% PFA and stained with anti-K10 antibodies (Abcam, ab9026). Nuclei were stained with DAPI. Red: Cdt1-mKO2 (G1 phase); Green: Geminin-mAG (S/G2/M phase); Blue: DAPI; Cyan: K10. (Original magnification, x20. Scale bar=200μm) (C-D) FUCCI NIKS cells transduced with retroviral vector harbouring EV, 11E6 or 16E6 were plated at 12 different densities in 96 well plates. After 72 hours, NIKS cells were fixed and stained with anti-K10 antibodies. Nuclei were stained with DAPI. Images for 9 fields in each well were captured by High content confocal imaging. The number of cells, % Geminin positive cells and % K10 positive cells in each field were quantified with Harmony image analysis software (Perkinelmer). The area of each field is 0.42mm^2^.

### E6AP and NHERF1 are crucial for E6 functions in cell proliferation and differentiation

After demonstrating that the E6 modifies the proliferation-differentiation trigger point, we then explored the underlying mechanism by focusing on existing E6 target proteins. E6AP is a conserved binding partner for Alpha group E6 proteins, we hypothesised that E6AP has a role in E6-regulated phenotypes as described above. In addition, NHERF1 was one of the first cellular proteins discovered that can be degraded by both high and low-risk E6 proteins (Accardi et al., 2011; Id et al., 2019). It interacts with a range of signalling proteins including YAP, PTEN and frizzled receptors (Georgescu et al., 2016; Wheeler et al., 2011). Several previously described E6AP-binding deficient mutants (Nicole Brimer, Charles Lyons, 2007; Oh et al., 2004; Zimmermann et al., 1999), 11E6^W133R^, 11E6^L111Q^ and 16E6^L50G^, were constructed (table 1). NHERF1-binding deficient 16E6^F69A^ mutant was established based on the work of Drews *et al*., 2019 and a corresponding point mutation in 11E6 was also generated (11E6^L70A^). Both mutants were validated for NHERF1 degradation (figure 2-supplementary 1) and included in the same experiment with FUCCI-expressing NIKS cells. Intriguingly, the ability of E6 to modulate proliferation, differentiation and saturation density was compromised by losing E6AP binding. As shown in figure 2B, cells expressing 11E6^W133R^ and 11E6^L111Q^ did not reach the same saturation density as the cells expressing 11E6^WT^. The percentage of geminin-positive cells was also decreased at post-confluence and cells started to differentiate at a lower density compared to NIKS expressing 11E6^WT^. Similarly, L70A mutation also abolished 11E6’s ability in regulating cell cycle, density and differentiation as well, suggesting an important role for NHERF1 degradation in 11E6 functions. For 16E6 group shown in figure 2C, all three mentioned phenotypes for NIKS cells expressing 16E6^L50G^ were lost, whereas 16E6^F69A^-expressing NIKS cells resembled the behaviour of 16E6^WT^ NIKS. This indicates E6AP is a significant contributor of 16E6 functions in the regulation of proliferation-differentiation switch. Furthermore, it appears that NHERF1 degradation plays a more critical role in this process for 11E6. While E6AP is a direct target of Alpha group E6, NHERF1 is one of the several secondary targets of the 16E6-E6AP complex. It is possible that other identified targets of 16E6 such as the DLG, scribble and other PDZ proteins are involved in this process (Vats et al., 2019).

**Table 1:** summary of E6 mutants used in this study. E6 mutants utilised in this work and associated functional defects.

| E6 mutant | Functional consequence | Reference |
| --- | --- | --- |
| 11E6<sup>L70A</sup> | Loss of NHERF1 degradation | Validated in figure 2 supplementary 1 |
| 11E6<sup>L111Q</sup> | Loss of E6AP binding | Oh et al., 2004; Brimer et al., 2007 |
| 11E6<sup>W133R</sup> | Loss of E6AP binding | Oh et al., 2004; Brimer et al., 2007 |
| 16E6<sup>L50G</sup> | Loss of E6AP binding | Zimmermann, Holger, et al., 1999 |
| 16E6<sup>F69A</sup> | Loss of NHERF1 binding/degradation | Drews et al., 2019 |

**Figure 2.**
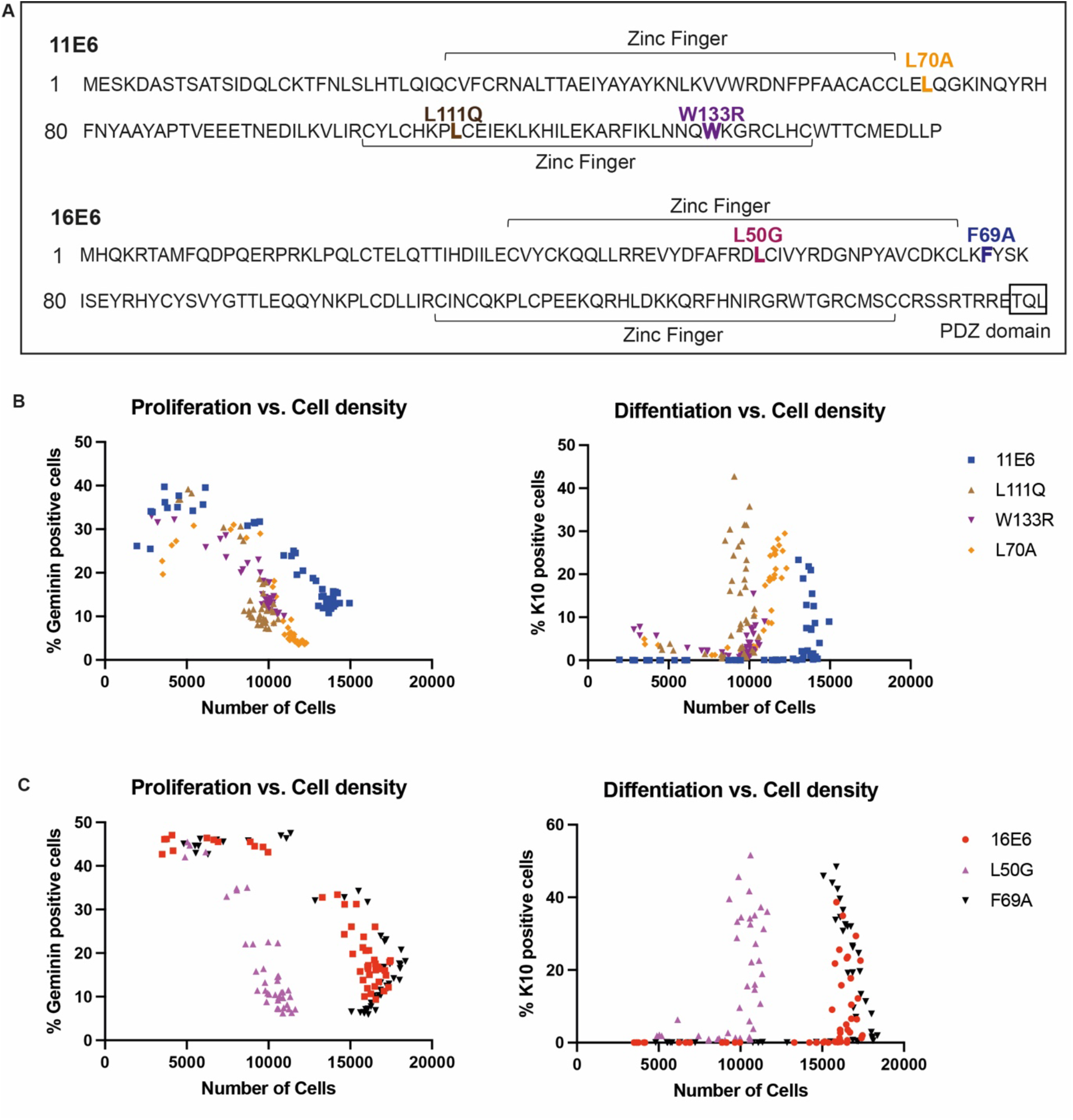
Role of E6AP and NHERF1 in E6-regulated keratinocyte phenotypes. (A) Amino acid sequences of 11E6 and 16E6. Point mutations are indicated by coloured texted. (B) 11E6 requires E6AP binding or NHERF1 binding to modulate proliferation, differentiation and saturation density of NIKS cells. FUCCI NIKS transduced with retroviral vectors harbouring 11E6 or 11E6 mutants were cultured to grow at 12 different cell densities. Cells were fixed with 4% PFA and stained with anti-K10 antibodies. Nuclei were stained with DAPI. % Geminin positive cells, % K10-positive cells and the number of cells were quantified in each field. The area of one field is 0.42mm^2^. (C) 16E6 requires E6AP but not NHERF1 binding to modulate proliferation, differentiation and saturation density of NIKS cells.

### E6AP but not NHERF1 contributes to E6’s competitive advantage in the lower layer of keratinocytes

Given E6’s functions identified above, we then sought to explore the consequences at the cell population level. Cell-cell competition assays enable us to study not only the ‘enhanced fitness’ that E6 confers on keratinocytes in the lower layer, but also allow us to assess keratinocyte delamination. Competition assays described in Saunders-Wood *et al*., 2022 was used as a model to mimic aspects of epithelium basal layer. NIKS^mCherry^ cells transduced with either E6^WT^ or E6 mutants and NIKS^eGFP^-EV cells were seeded at the same ratio to form a confluent monolayer on day one. Over a course of nine days, the cell populations were grown to form at least two layers and fixed at each time point. Images of the lower layer and upper layer of cells were captured by Z-stack confocal microscopy (figure 3A). Day nine images are presented to show the most obvious effect. On day one, all experimental groups started at 50:50 ratio for mCherry and eGFP NIKS cells. However, inhibitory effect on cell growth was observed for mCherry cells in comparison to eGFP cells, thus the proportion of NIKS^mCherry^-EV cells against NIKS^eGFP^-EV cells dropped below 50% at day 7 and day 9 (figure 3B). Despite this, the 11E6- and 16E6-expressing NIKS^mCherry^ cells gradually increased in proportion and outcompeted NIKS^eGFP^-EV cells, reaching 71.6% and 83.3% coverage respectively in the lower layer on day nine. This suggests that the NIKS^eGFP^-EV cells were ‘less-fit’ as they were displaced by NIKS^mCherry^-E6 cells from the lower layer and moved to the upper layer (figure 3A). Furthermore, E6AP-binding mutant and NHERF1-binding mutant cell lines enabled us to investigate the contribution of E6AP and NHERF1 in E6 functions during cell-cell competition (Figure 3C-D). We found that the ability of both cell populations expressing 11E6 E6AP-binding mutants to persist in the lower layer was significantly compromised in comparison to 11E6^WT^. On day nine, both NIKS^mCherry^-11E6^W133R^ and NIKS^mCherry^-11E6^L111Q^ cells reached about 60% in the lower layer, retaining a slight growth advantage against NIKS^eGFP^ cells. They also lost the ability to displace NIKS^eGFP^ cells into the upper layer (figure A). However, NHERF1-binding mutant 11E6^L70A^-expressing NIKS^mCherry^ cells behaved in a similar manner as NIKS^mCherry^-11E6^WT^, suggesting NHERF1 may not be involved in E6 function during competition assay. In parallel, we noticed that the competitive advantage of 16E6^L50G^ mutant-expressing NIKS^mCherry^ cells was also significantly compromised in comparison to NIKS^mCherry^-16E6^WT^ and occupied approximately 67.9% in the lower layer. 16E6^F69A^ mutant-expressing NIKS^mCherry^ cells resembled the trend of NIKS^mCherry^-16E6^WT^ cells at all time points, reaching about 79% at day 9. This further indicates that NHERF1 may not play an important role in E6 regulation of cell delamination. This is also consistent with recently published results (Brimer & Vande Pol, 2022).

**Figure 3.**
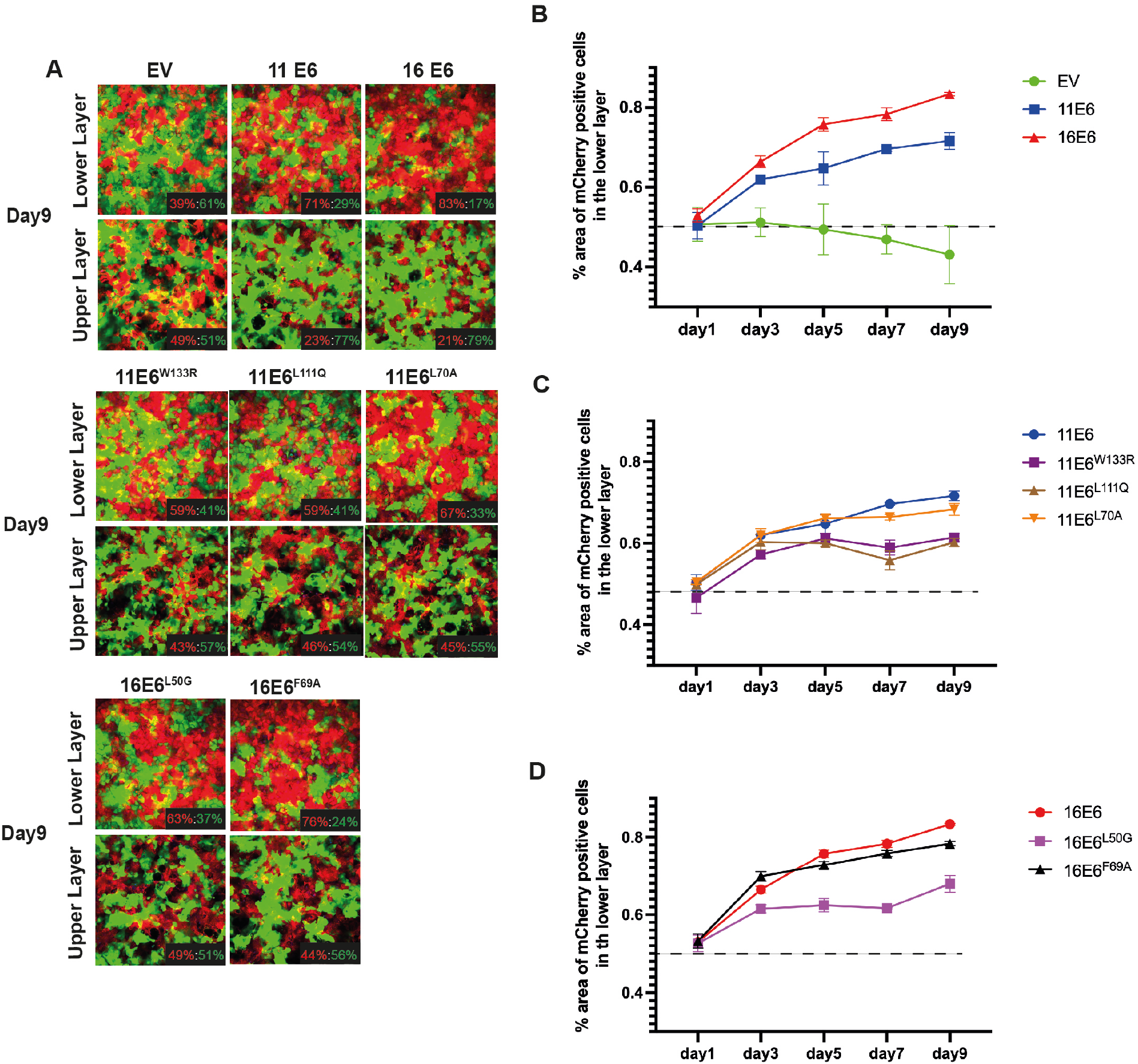
NIKS cells expressing E6 preferentially persist in the lower layer of cells in a high-density competition assay. (A) Representative images of the lower and upper layers of each group on day nine are shown: NIKS-eGFP-EV cells were seeded together with NIKS-mCherry-EV or NIKS-mCherry cells expressing 11E6/11E6^W133R^/11E6^L111Q^/11E6^L70A^/16E6/16E6^L50G^/16E6^F69A^. Ratio of area occupied by mCherry cells to eGFP cells is presented at the lower right corner of each image. Images were captured and quantified by Harmony high content imaging and analysis software (Perkinelmer). Original magnification: x20. (B-D) Graphs showing the % area of NIKS mCherry cells in the lower layer over the course of competition assay. NIKS mCherry for each cell line against NIKS eGFP was seeded at ratio 50% : 50% and were cultured for nine days. The plates were then fixed by 4% PFA at day 1, 3, 5, 7 and 9, followed by staining with DAPI. The lower layer of cells was scanned by Confocal microscopy. The area of mCherry cells was quantified for each group of cell lines and data are means ± standard errors of three random fields. The area of one field is 0.42mm^2^.

### Condyloma staining revealed several cellular targets regulated during low-risk HPV productive lifecycle

Condyloma acuminatum is HPV-induced squamous epithelial proliferation in the anogenital region, caused by low-risk HPV types 6 and 11 (Gupta et al., 2018). Figure 4A shows a condyloma biopsy collected from patients infected with HPV11 and stained with haemotoxylin & eosin (H&E). It comprises of both lesion and uninfected area, allowing us to make direct comparisons with the following biomarker analysis. To confirm the expression of HPV viral genes in epithelium basal layer, RNAscope was performed on condyloma biopsies to indicate the expression pattern of HPV11E6E7 mRNA (figure 4B). 11E6E7 mRNA expression was restrained in the basal layer and the lower layers of the lesion. The mRNA abundance was increased significantly in the middle and upper layers. This is typical during low-risk HPV productive infection. In the same region, biomarkers such as MCM7 and K10 were applied to provide indications for cell proliferation and differentiation status (figure 4C-D). Quantification of nucleus per μm shows an elevated cell density in the lesion than uninfected epithelium (figure 4E). Additionally, percentage of MCM7 positive cells was significantly increased in the lesion, suggesting enhanced cell proliferation (figure 4F). The increased distance from basal lamina to the cells starting to express K10 indicated a delay in the commitment to differentiation (figure 4G). This suggests that the timing of differentiation marker expression was delayed in infected cells. Overall, these results are in correlation with our findings in phenotypic assays (figures 1 and 2).

**Figure 4.**
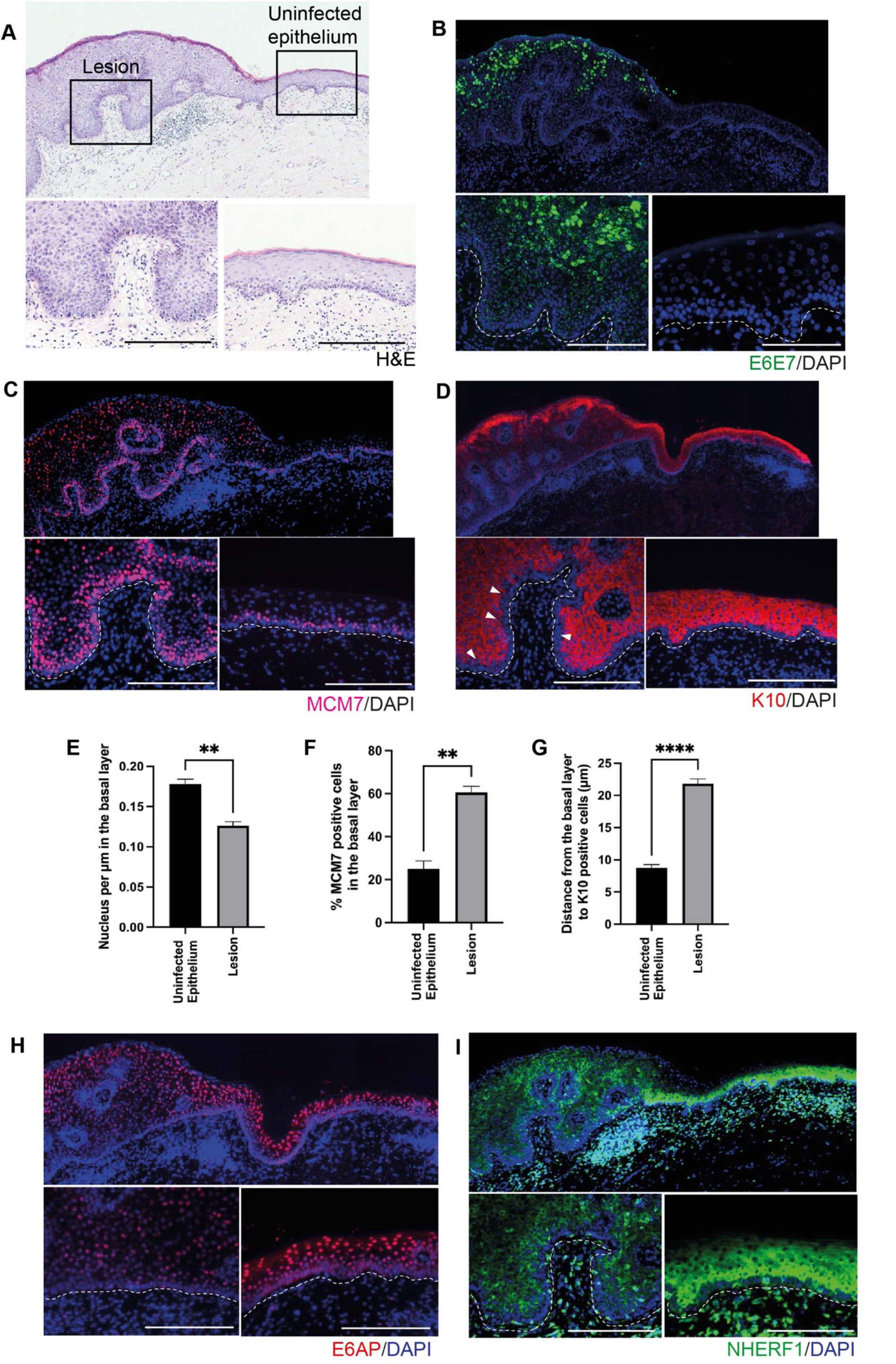
Localisation and expression pattern of E6/E7 and its targets during HPV11 productive lifecycle in condyloma acuminatum. (A) H&E staining of condyloma acuminatum biopsy with enlargement areas of lesion (left) and non-infected epithelium (right). (B-D) E6/E7 RNAScope, MCM7 and K10 immunofluorescence staining was carried out on adjacent sections. Nuclei were counterstained with DAPI (blue). The scale is shown with a white bar (200 μm). Enlargement images are shown at the bottom, lower left is the lesion and lower right is the non-infected area. The dotted lines indicate the position of the basal layer. (E-G) The graphs show quantification of cell density (nucleus per μm) (E), % MCM positive cells in the basal layer (F) and distance between the basal lamina to the bottom of k10 positive cells (μm) are presented in column graphs. Quantification is done by manully counting cells with Image J. Graphs show mean values ± standard errors of at least three fields from condyloma biopsies. P values were calculated with student t tests. **, *P* ≤0.01; ****, *P* ≤0.0001. (H-I) E6AP and NHERF1 protein expression in infected (lower left) and non-infected (lower right) region of the biopsy. The scale is shown with a white bar (200 μm).

To assess the clinical relevance of our observations on cell culture, E6AP and NHERF1 staining were applied on adjacent sections of the several condyloma biopsies. The localisation and expression pattern of these proteins were examined in both infected and non-infected tissue areas. Few previous studies reported the expression pattern of E6AP in human stratified epithelium. However, we found that in non-infected area, E6AP was predominantly cytoplasmic in the basal layer (figure 4H). From the second layer and the above, E6AP nuclear localization became progressively more evident. In the lesion area where the papilla starts to present, there was significant reduction of nuclear E6AP in the upper layers. Additionally, the cytoplasmic E6AP in the basal layer was decreased in comparison to the non-infected area. Overall, there was a general reduction in E6AP protein abundance in the lesion. This agrees with previous findings that E6 can induce the auto-ubiquitination of E6AP and its degradation (Kao et al., 2000). Further, our *in vitro* studies showed that NIKS cells stably expressing either 11E6 or 16E6 had lower E6AP protein levels compared to control cells (figure 4-supplementary 1A). In 3D organotypic rafts, reduction of E6AP was identified throughout the bottom and upper layers (figure 4-supplementary1B). In addition, NHERF1 was mostly expressed from the second layer and upper layers in non-infected area, which displayed a cytoplasmic and perinuclear pattern (figure 4I). There were a few basal cells found expressing NHERF1. However, NHERF1 levels were remarkably decreased in the lesion, supporting previous literature that E6 degrades NHERF1 in various cell lines (Accardi et al., 2011;Drews et al., 2019).

### E6 regulates YAP localization and phosphorylation level via E6AP and NHERF1

YAP has been identified as a critical modulator in sensing cell density to regulate cell proliferation, and has been shown to have important functional roles in keratinocyte homeostasis (Corley et al., 2018; Elbediwy et al., 2016; Zhao et al., 2007). YAP localization is mainly regulated through phosphorylation by LATS1/2 (Bernascone & Martin-Belmonte, 2013). At high cell density, a major phosphorylation of YAP occurs on the position Serine 127 (Ser127), leading to YAP sequestration in the cytoplasm (Zhao et al., 2010). At low cell density, YAP is not phosphorylated and enters the nucleus to activate downstream genes (M. K. Kim et al., 2018). Past work has demonstrated that high-risk HPV E6 proteins regulate the Hippo signalling cascade during the progression to cervical cancer (He et al., 2015). However, we believe both low-risk and high-risk E6 proteins manipulate YAP activity in low-grade lesions to adjust homeostasis. Therefore, we examined the localisation and abundance of YAP in the condyloma tissue by using antibody that specifically recognises the non-phosphorylated (active) form of YAP1 (figure 5A). In non-infected epithelium, YAP is predominately nuclear when it is present in the basal cells. This is consistent with the observations from previous reports (Elbediwy et al., 2016; Vincent-Mistiaen et al., 2018; Xiao et al., 2014). Nuclear YAP gradually decreased in the suprabasal and upper layers and became more prominent in the granular or cornified layer. By comparison, the number of cells with prominent level of YAP increased in the basal layer in the infected area, suggesting a subtle modulation of homeostasis towards proliferation. Ser127 YAP was found significantly decreased in the basal and suprabasal layers, whereas relatively high-level expression of ser127 YAP is observed in non-infected tissue (figure 5B). This implicates HPV viral gene expression in the basal layer modulates YAP nuclear-cytoplasmic shift.

**Figure 5.**
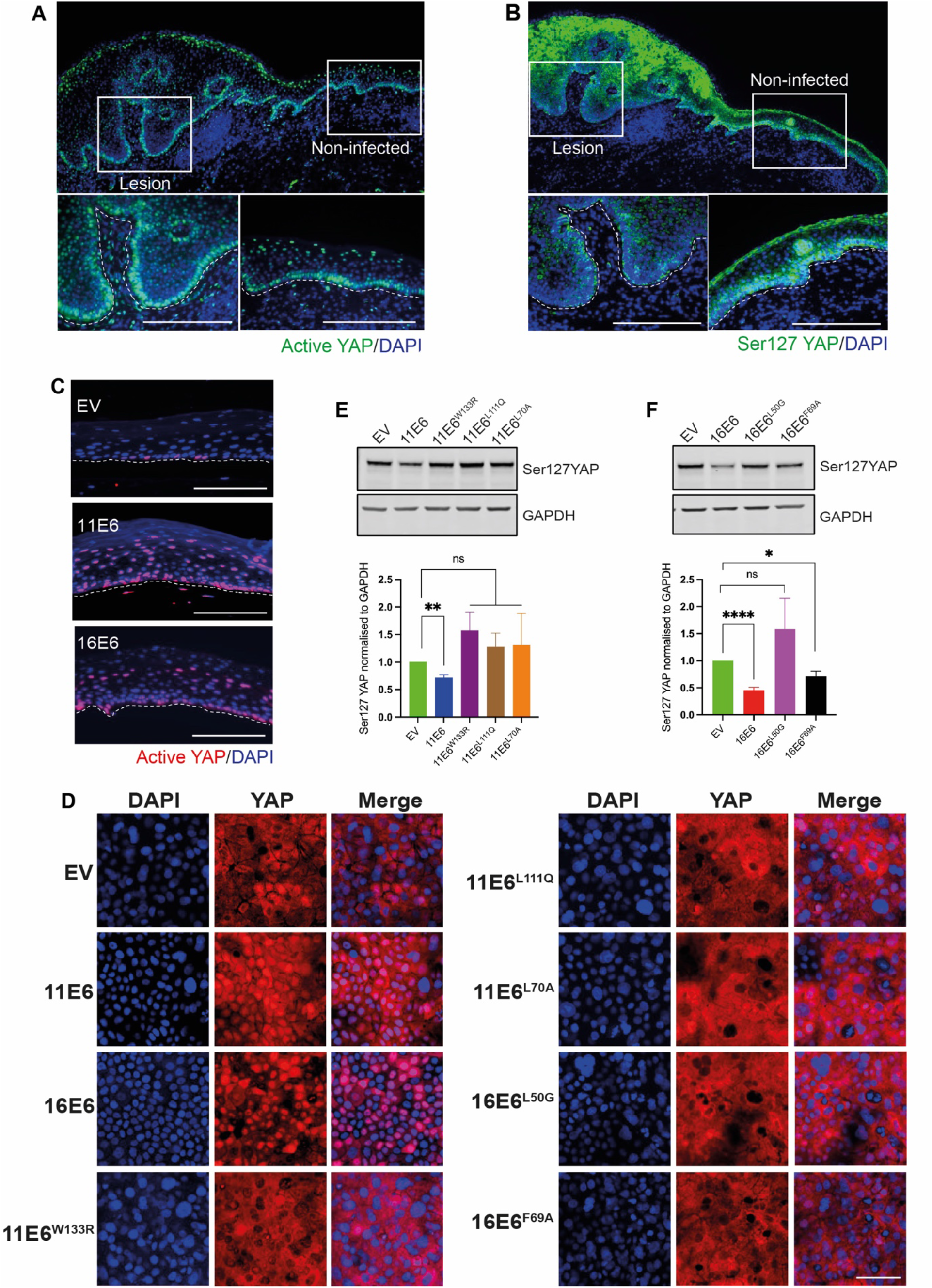
HPV E6 requires E6AP and NHERF1 to enhance YAP nuclear localisation. (A-B) Active YAP antibody (Abcam, ab205270) that recognizes un-phosphorylated form of YAP and p-YAP (Ser127) antibody (Cell signalling, 4911) that only recognizes YAP phosphorylated at position Serine 127 were stained on condyloma biopsy. Nuclei were stained with DAPI. Specific areas of lesions and non-infected are shown as enlargement images at the bottom, lower left is the lesion and lower right is the non-infected epithelium. (C) Organotypic rafts of NIKS transduced with retroviral vectors encoding EV, 11E6 and 16E6 were established, sectioned and stained with active YAP antibody. The scale for all images is shown with a white bar (200 μm). (D) NIKS transduced with retroviral vectors encoding EV, 11E6, 16E6, 11E6^W133R^, 11E6^L111Q^, 11E6^L70A^, 16E6^L50G^, 16E6^F69A^ were fixed at post-confluence, followed by staining with active YAP antibody and DAPI. Scale = 100 μm. (E-F) NIKS cells at post-confluence were collected and whole-cell lysates were subjected to Western blot analysis for P-YAP (Ser127). In all quantified Western blotting results, representative blots are shown. Data are means ± standard errors of three biological replicates. ****P < 0.0001, **P < 0.01 (two-tailed Student’s t test), ns, not significant.

In parallel, NIKS transduced with retroviral expression vectors encoding either 11E6 or 16E6 were used to grow organotypic cultures, which displayed higher level of nuclear YAP in the basal layer of the rafts relative to parental NIKS raft (figure 5C), supporting our observations in condyloma tissues. To study how E6 overcomes the impact of contact inhibition through YAP activation, we seeded FUCCI NIKS cells expressing E6 or E6 mutants at post-confluence and stained with active YAP. 11E6 and 16E6 both led to enhanced nuclear YAP relative to NIKS-EV, whereas the E6 mutants 11E6^W133R^, 11E6^L111Q^, 11E6^L70A^, 16E6^L50G^ and 16E6^F69A^ lost the ability to retain YAP in the nucleus (figure 5D, figure 5-supplementary1). This suggests that E6AP and NHERF1 are involved in YAP nuclear localisation. Also, E6-expressing NIKS cells had reduced Ser127 YAP levels at post-confluence, whereas the mutant cell lines had similar levels of Ser127 YAP as NIKS-EV (figure 5E-F). This further implies that E6 requires NHERF1 and E6AP binding to promote YAP nuclear localisation at post-confluence, and this leads to the reduction of phosphorylated YAP in cytoplasm.

### E6AP is important for cell differentiation

Based on the results shown above, our work demonstrates that E6AP plays an important role in E6-regulated homeostatic phenotypes in keratinocytes. Accumulating evidence shows that Alpha group E6 binding to E6AP leads to the activation of its ubiquitin ligase activity and degradation(Brimer et al., 2017; Kao et al., 2000). Together with our results, it prompts the hypothesis that E6 may regulate the natural cellular targets of E6AP through directly targeting E6AP for degradation. This prompted us to generate NIKS cells transduced with shRNA oligonucleotides targeting E6AP. The knockdown effect was validated with western blot (figure 6A). NIKS-shRNA-luciferase (control) and NIKS-shRNA-E6AP cells were plated at cell densities ranging from pre-confluence to post-confluence. After 72 hours, cells were fixed and stained with K10. NIKS-shRNA-E6AP cells displayed higher saturation density (13,000 cells/field) and delay of K10 expression in comparison to the control NIKS (figure 6A). At the same cell density, NIKS cells positive in the proliferation marker MCM7 cells were increased, whereas NIKS cells expressing K10 were significantly reduced (figure 6B-C). This provides evidence that E6AP contributes to the balance of proliferation-differentiation switch in keratinocytes. Further, NIKS organotypic rafts with E6AP knocked down were established, which allowed us to examine K10 expression in different layers. We found a slight delay of k10 expression in the second layer of the raft expressing shRNA E6AP (figure 6-supplementary 1).

**Figure 6.**
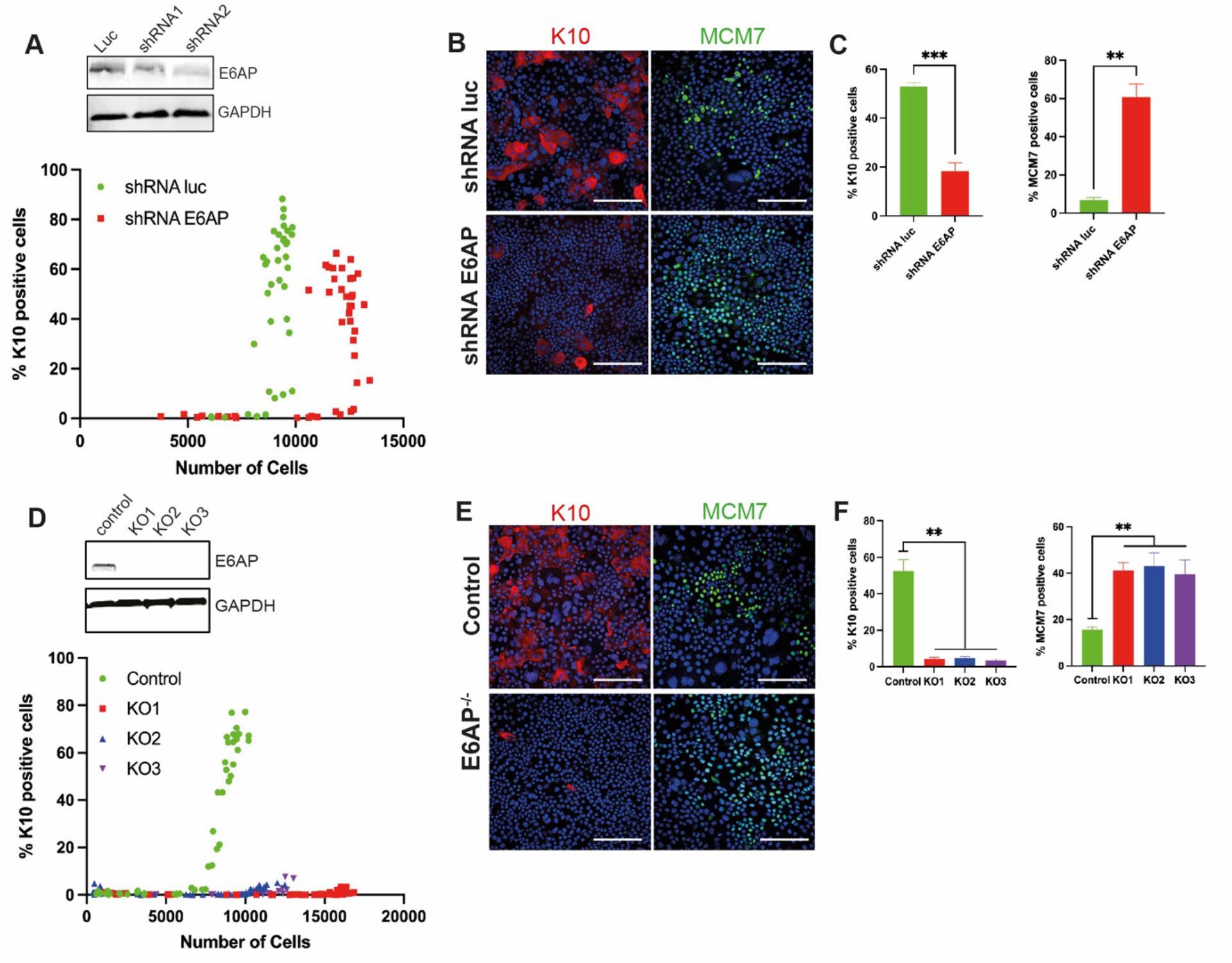
E6AP is a critical regulator of keratinocyte differentiation. (A) E6AP was knocked out in NIKS cells by transfecting cells with px459 plasmid expressing E6AP gRNA. Puromycin and single cell selection was carried out to obtain three independent KO clones. The control cells were NIKS cells transfected with px459 plasmid expressing gRNA targeting a non-existing gene in mammalian cells. Western blot indicates the loss of E6AP in NIKS^E6AP-/-^ cells. Control and knockout cell lines were plated at different densities and fixed after 72 hours. Cells were then stained with K10 and DAPI. % K10 positive cells was plotted against the number of cells per field. (B-C) Control and NIKS^E6AP-/-^ cells at post-confluence were stained with K10 (red) and MCM7 (green). % K10 positive cells and % MCM7 positive cells were quantified for at least three representative images, mean values ± standard errors are shown in the graphs. Student t-test was performed between each group. (D) NIKS cells were retrovirally transduced with plasmids expressing shRNA targeting E6AP or luciferase. Cells were then selected with puromycin and validated with western blot. Cells were plated at different densities and after 72 hours, cells were stained with K10 and DAPI. % K10 positive cells was plotted against the number of cells per field. (E-F) NIKS-shRNA-luc and NIKS-shRNA-E6AP cells at post-confluence were stained with K10 (red) and MCM7 (green). % K10 positive cells and % MCM7 positive cells were quantified for at least three representative images, mean values ± standard errors are shown in the graphs. Student t-test was performed between each group.

In parallel, NIKS cell lines with E6AP knocked out by CRISPR-Cas9 were established. Three clonal E6AP^-/-^ NIKS cell lines with genome edited by gRNA1 were selected. Sequencing of the genomic region targeted by the gRNA confirmed frameshift mutations had been introduced into each allele and no wild-type (WT) allele remained. Consistent with this, immunoblotting showed loss of E6AP expression (figure 6D). A control NIKS cell line expressing gRNA targeting a random rice gene was also established alongside and underwent single cell selection. The E6AP^-/-^ NIKS cell lines were then plated in cell densities ranging from pre-confluence to post-confluence. After 72 hours, cells were fixed and stained with K10 (figure 6E). Three E6AP^-/-^ NIKS cell lines all reached higher saturation densities, from 12,000 cells/field to 17,000 cells/field in comparison to the control cell line (8000 cells/field). K10 expression was remarkably reduced in E6AP^-/-^ NIKS cells after saturation density was reached. Comparing at similar cell densities, K10 expression was significantly lower and %MCM7 positive cells was increased (figure E-F). These results support our observations in NIKS-shRNA-E6AP cells and demonstrate the potential role of E6AP in epithelial homeostasis.

### E6 requires E6AP depletion to impair differentiation gene expression and activate YAP target genes in keratinocytes

To establish which cellular pathways are affected following E6AP loss in keratinocytes under condition which differentiation would normally be triggered, total RNA was isolated from three independent samples of NIKS-control and NIKS E6AP^-/-^ cells that grew until post-confluence. With the standard mRNA read depth of around 20 million reads/sample, 3824 genes were differentially expressed with fold-change >=2 and adjusted P<=0.05 in E6AP^-/-^ cells (figure 7-supplementary 1A). Of these, 1664 genes were down-regulated and 2160 genes were upregulated in the absence of E6AP. In the gene enrichment analysis, more than half of the down-regulated genes were in keratinocyte differentiation-related GO categories (figure 7A and B). These included cornification, keratinisation, epidermis development, keratinocyte differentiation, skin development and epidermal cell differentiation. We selected a subset of markers of keratinocyte differentiation, such as keratin 1, keratin 4, keratin 10, keratin16 and involucrin for validation by qRT-PCR. These genes were all significantly downregulated in E6AP^-/-^ cells (figure 7C). KRT1, KRT4, KRT10, and KRT16 are cytokeratins associated with the suprabasal layers of differentiating keratinocytes (Sharma et al., 2019; Werner et al., 2020). Involucrin (IVL) is also a marker for keratinocyte differentiation commitment which expresses at high levels in the suprabasal layers of the epidermis before cornification occurs (Sanchez-Danes & Blanpain, 2018). Additionally, relevant GO categories upregulated by E6AP^-/-^ included regulation of signalling receptor activity, extracellular matrix organisation and cell-cell adhesion etc (figure 7-supplementary 1C). Among these GO terms, we found a subset of YAP target genes were significantly upregulated (figure 7B). qPCR was then performed to quantify to verify the relative abundance of YAP downstream gene expression. Consistent with the RNA-seq outcome, well-characterised YAP target genes *AREG, PLAU, PTGS2, AXL* and *CTGF* that were described in previous literature (Corley et al., 2018; Franklin et al., 2020; H. Kim et al., 2021; Li et al., 2020), were expressed at 2-to 20-fold higher levels when E6AP was depleted (figure 7C). All these genes have indicated functions in driving cell proliferation.

**Figure 7.**
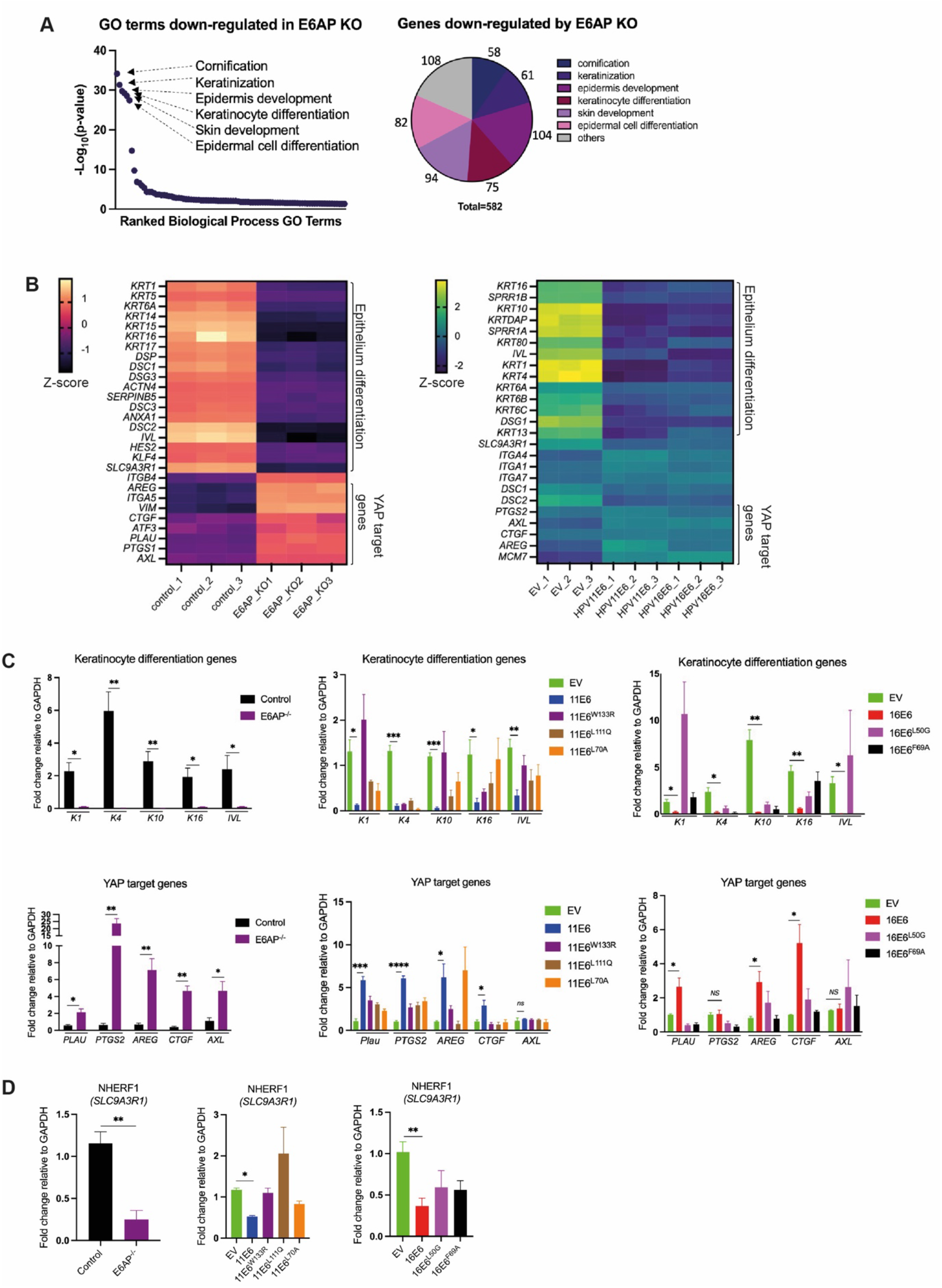
E6 requires E6AP depletion to impair differentiation gene expression and activate YAP target genes in keratinocytes. Total RNA was extracted from NIKS control, NIKS E6AP^-/-^ and NIKS transduced with EV, 11E6 or 16E6 when the cells reached post-confluence. PolyA selected RNA was analyzed by RNA-seq. (A) GO enrichment analysis of genes down-regulated in NIKS E6AP-/-compared with NIKS-control. Pie chart displays the fraction of genes down-regulated in the absence E6AP that fall into enriched GO Terms. (B) FPKM (expected number of Fragments Per Kilobase of transcript sequence per Millions base pairs sequenced) values were calculated to estimate gene expression levels from RNA-seq results. Selected epithelium differentiation genes and YAP-responsive genes that was significantly differentially expressed are shown in the heatmap. The legend shows the range of Log_2_ (FPKM+1) values of genes that are homogenised across the row (Z-score). Complete set of heatmap is shown in supplementary file 2. (C) Transcript abundance for keratinocyte differentiation genes or YAP target genes was measured in NIKS-control, NIKS E6AP^-/-^, NIKS-EV, NIKS-11E6, NIKS-11E6^W133R^, NIKS-11E6^L111Q^, NIKS-11E6^L70A^, NIKS-16E6, NIKS-16E6^L50G^, NIKS-16E6^F69A^ by qRT-PCR. Bar graphs display mean +-standard errors of three independent experiments. ***P < 0.001, **P < 0.01 (two-tailed Student’s t test), *P < 0.05, ns, not significant. (D) NHERF1 (*SLC9A3R1*) mRNA level was quantified by qPCR in three independent experiments.

Given our finding that E6 requires E6AP binding to promote YAP nuclear localisation (figure 5), we hypothesised that E6 inhibits E6AP function to activate specific YAP target genes. Total RNA was extracted from NIKS-control and NIKS cells stably expressing either 11E6 or 16E6 that grew until post-confluence to trigger differentiation, and sent for RNA sequencing. For the GO enrichment analysis, both 11E6 and 16E6 downregulated differentiation-related processes including cornification, keratinisation, keratinocyte differentiation etc (figure 7 supplementary 1C). At the same time, cell cycle-related processes including G2/M phase transition, G1/S phase transition and cell division were upregulated (figure 7 supplementary 1C). These results correlate to the increased proliferation and delayed differentiation of NIKS cells under the expression of E6 presented above (figure 1). All the YAP target genes upregulated in E6AP^-/-^ cells were also found to be activated in E6-expressing NIKS cells, including *PTGS2, AXL, CYR61, CTGF* and *AREG* (figure 7C). To confirm that E6AP degradation is required for YAP target gene activation, NIKS cells expressing E6 or the mutants were seeded at high cell density and the mRNA abundance was quantified by RT-qPCR. Indeed, 11E6^W133R^, 11E6^L111Q^ and 16E6^L50G^ which cannot induce E6AP degradation had reduced ability to upregulate YAP-responsive genes (figure 7C). Interestingly, RNA-seq and qPCR results suggest that NHERF1 gene (*SLC9A3R1*) downregulation is found in E6-expressing and E6AP^-/-^ NIKS cells (figure 7D). Thus, NHERF1 is not only degraded by E6-E6AP complex but also downregulated by E6 at transcriptional level. It is possible that NHERF1 reduction directly controls YAP transcriptional activity, because E6 mutants 11E6^L70A^ and 16E6^F69A^ cannot upregulate YAP target genes.

## Discussion

Human papillomaviruses establish chronic lesions in the epithelium (Doorbar et al., 2012; Stanley, 2012). During evolution, HPVs became adapted to niches through manipulating molecular processes and hence developed distinct tissue tropisms (Doorbar et al., 2021; Kranjec & Doorbar, 2016). For example, the Alpha genus E6 preferentially bind to E6AP whereas E6 from the other genera confer stronger interaction with MAML (Brimer et al., 2012, 2017; Tan et al., 2012). In both instances, this leads to the inhibition of Notch signalling and delay of terminal differentiation program (Kranjec et al., 2017; Meyers et al., 2017; Tan et al., 2012). Recent published work on MmuPV and high-risk E6 has indicated that E6 rather than E7 has a dominant role in inducing cell competition and promotes basal cell retention (Brimer & Vande Pol, 2022; Saunders-Wood et al., 2022). Also, our previous work have demonstrated that low-risk E6 is the main driver for keratinocyte proliferation, and it plays a major role in delaying keratinocyte committing to differentiation (Kranjec et al., 2017; Murakami et al., 2019). Therefore, accumulating evidence suggests that E6 proteins have conserved functions in modulating the balance between cell proliferation and differentiation that are crucial for HPV-infected lesion expansion and persistence. Thus, our work firstly dissected the shared functions between high- and low-risk HPV E6 and found that they target similar homeostatic processes, in both cases E6AP serves an important role.

In this study, our results clearly show that high or low-risk E6 expression increased the proportion of cycling cells at both pre-confluence and post-confluence (figure 1). Because E6 has a prominent role in overcoming normal keratinocyte contact inhibition (Kranjec et al., 2017; Luna et al., 2021; Zheng et al., 2022), cells expressing E6 typically reached a higher saturation density than the control cells. Nevertheless, 16E6-expressing cells always reach higher saturation density whereas 11E6-expressing cells reach lower density. Similarly, NIKS transduced with either E6 typically start to show K10 expression at higher cell density, with 11E6 causing the intermediate phenotype between the control and 16E6. With cell-cell competition assays, we examined the progression of E6-expressing cell phenotype from the first layer to the second layer. This is in line with current thinking that E6 expression retains keratinocyte in the bottom layer and delays delamination, while the wild type NIKS cells were displaced and entered the second layer. Although 11E6 appeared to have the intermediate phenotype, they share a basic set of functions to modulate cellular phenotypes involved in homeostasis. In both cases, E6AP plays an important role. The more subtle cellular phenotypes caused by 11E6 expression comparing to 16E6 could be partially due to their different modes of interaction with E6AP. It was suggested that 11E6 and 16E6 bind to various auxiliary regions on E6AP that lead to distinct substrate degradation (Drews et al., 2020). For example, 11E6 cannot cause p53 degradation but degrades NHERF1 in a similar way as 16E6. Also, in both previous *in vitro* binding or co-immunoprecipitation studies, interaction between 11E6 and E6AP has shown to be weaker than 16E6-E6AP binding (Brimer et al., 2007; Cooper et al., 2003).

Currently, Hippo signalling has emerged as one of the key pathways being altered frequently in HPV-related cancers (Olmedo-Nieva et al., 2020). It has been suggested that high-risk E6 requires the PDZ motif to promote YAP nuclear localisation in serum-starved keratinocytes (Webb Strickland et al., 2018). High-risk E6 was also shown to drive cervical cancer cell proliferation by maintaining high levels of YAP in cells and YAP expression is correlated with cervical cancer progression (He et al., 2015). More recently, high risk HPV E7 was proposed to activate YAP1 in basal keratinocytes by degrading PTPN14, which contributes to papillomavirus persistence and carcinogenesis (Hatterschide et al., 2019, 2022). Despite its critical involvement in carcinogenesis, YAP is also required for normal skin homeostasis (Akladios et al., 2017; Georgescu et al., 2016). Proliferation of basal layer cells was significantly reduced in YAP/TAZ double knockout mouse skin (Elbediwy et al., 2016). Our finding revealed that YAP1 nuclear translocation can be promoted by low-risk 11E6 as well, and this is achieved through E6AP binding, suggesting YAP1 activation is involved during both low-risk and high-risk HPV infections (figure 5). Highly conserved feature of E6 binding to E6AP indicates that YAP1 activation and maintenance of basal cell state is likely shared among diverse Alpha genus E6 proteins. RNA sequencing performed on basal layer human keratinocytes indicate that YAP transcriptional regulation is active exclusively in the basal cell population (Elbediwy et al., 2016). This correlates with our observation that active YAP expression is mostly identified in the basal layer of condyloma section and upregulated in the presence of low-risk HPV infection (figure 5). Further, RNA sequencing and qPCR validation demonstrated that YAP downstream genes were activated in the presence of either 11E6 or 16E6. Loss of E6AP binding abolished E6 function in YAP downstream gene upregulation (figure 7). Our results are consistent with previous findings that *PLAU* and *PTGS2* are positively regulated by constitutive YAP activity in proliferating keratinocytes in the mouse skin in vivo, and in HaCat keratinocytes grown in vitro (Corley et al., 2018). Thus, both E6 drive keratinocyte proliferation through activating YAP downstream genes. It was recently shown that YAP/TAZ regulates differentiation genes in keratinocytes in the basal layer of organotypic raft culture (Hatterschide et al., 2022). In addition, YAP activation crosstalk with the Notch signalling by upregulating DLL1, JAG2 and DLL3 ligands, leading to cell-autonomous cis-inhibition of Notch (Totaro, Panciera, et al., 2018). This allows epidermal progenitors to maintain in an undifferentiated state. All of these ligands were found upregulated in our RNA-seq results for E6-expressing keratinocytes. Thus, the local microenvironment is dynamic regulated by E6 to orchestrate spatial control of self-renewal verses differentiation of basal layer progenitor cells.

Alpha genus E6 proteins deplete E6AP to different extent by inducing the self-ubiquitination and degradation of E6AP (Brimer et al., 2017; Kao et al., 2000). Importantly, our clinical observations show that in the basal layer where E6/E7 viral genes are expressed, a reduction of cytoplasmic E6AP was observed in comparison to uninfected epithelium. In the upper layers, although E6AP accumulates in the nucleus, its abundance is noticeably less prominent in the presence of amplified E6/E7 expression (figure 4). This is the first description of E6AP pattern in human tissue, which agrees well with our *in vitro* work that NIKS cells transduced with E6 led to decreased endogenous E6AP protein level (figure 4 supplementary 1). However, the consequence of E6AP degradation in the context of HPV life cycle and epithelium homeostasis has not been fully understood. Our results showed that loss of E6AP binding diminished a major component of both high and low-risk E6 functions in driving cell cycle entry, delaying differentiation, overcoming contact inhibition and basal cell retention (figure 2-3). Because high-risk E6 targets p53 to delay keratinocyte differentiation, low-risk E6 that cannot lead to p53 degradation may target E6AP directly. Our shRNA-E6AP and E6AP^-/-^ cell lines both demonstrated that E6AP is required for normal keratinocyte differentiation program and its depletion leads to less cells committing to differentiation and mostly remain in proliferative state. Additionally, past literature suggests that E6AP has impact on cell cycle control and proliferation (A Mishra & Jana, 2008; Amit Mishra et al., 2009; Srinivasan & Nawaz, 2011).

It is possible that E6 regulates the levels of natural cellular targets of E6AP through inducing its degradation. On the other hand, E6 modifies E6AP substrate specificity to degrade other cellular proteins can still contribute to the phenotypes we observed. Certainly, E6AP and NHERF1 are both depleted in cells expressing either 11E6 or 16E6. Also, NHERF1 expression level goes down in the absence of E6AP (figure 7). RNA-seq results showed the resemblance of E6-expressing and E6AP^-/-^ keratinocytes that both displayed lower expression of keratinocyte differentiation-related genes and higher level of YAP downstream genes. NHERF1 has been shown to directly interacts with YAP and its depletion leads to YAP translocation to the nucleus (Georgescu et al., 2016). Either E6 expression or E6AP knockout results in reduced NHERF1 mRNA expression in our RNA-seq and qPCR results (figure 7). Also, E6 mutants that cannot degrade NHERF1 failed to increase nuclear YAP (figure 5). This implicates that NHERF1 may also be critically involved in YAP activation.

As with the high-risk HPVs, low-risk HPVs are a group of evolutionarily successful viruses that are able to persist in epithelium basal layer. During lesion maintenance, the high-risk group can progress to neoplasia whereas the low-risk group cause significant mobility and healthcare burden (Saxena et al., 2022; Thapa et al., 2018). The current treatment with repeat surgical resection of papillomatous disease does not address the fundamental underlying issue of chronic infection with low-risk HPV and complete clearance of the reservoir of infected cells becomes more important (Egawa & Doorbar, 2017; R Ivancic et al., 2020; Ryan Ivancic et al., 2018). Despite the disease outcomes, low-risk and high-risk HPVs modulate similar pathways during lesion expansion and persistence. Successful establishment of persistent infection is a prerequisite for both low-risk HPV chronic lesion and high-risk HPV carcinogenesis (Doorbar et al., 2021). Our work discovered new ways of E6 interacting with cellular proteins to assist lesion maintenance, which shed light on potential therapeutic strategies such as small molecular inhibitors upon disease elimination.

## Materials and methods

### Cell culture

NIKS (a gift from Paul Lambert, McArdle Laboratory for Cancer Research, University of Wisconsin), a HPV-negative spontaneously immortalised human keratinocyte cell line, was maintained at sub-confluence on γ-Irradiated J2 3T3 feeder cells (a gift from Paul Lambert) in F medium with all supplements as previously described (Flores et al., 1999). 293T (ATCC) were maintained in Dulbecco’s Modified Eagle’s Medium (DMEM, SIGMA) supplemented with 10% fetal calf serum (FCS, HyClone) and 1% penicillin and streptomycin. FUCCI NIKS cells were established by transducing with the FUCCI cell cycle sensor and FACS sorted for high level expression of Cdt1-mKO2 (G1/G0 phase) and Geminin-mAG (S/G2/M phase). E6AP KO or mock control cell lines were established by transfection of px459 with sgRNA targeting E6AP or a non-exist gene (supplementary table 1).

### Plasmid construction and site-directed mutagenesis

pSpCas9(BB)-2A-Puro (PX459)-E6APgRNA plasmid and pSpCas9(BB)-2A-Puro (PX459)-rice gRNA plasmid were kind gifts from Lawrence banks (Jayashree Thatte, 2018) and Dr. Yongxu Lu from Department of Pathology, University of Cambridge. Construction of the retroviral vectors pQCXIN-Flag11E6 and pQCXIN-Flag16E6 were accomplished by cloning the coding sequence using Gateway Technology (Thermo Fisher Scientific, MA, USA) following manufacturer’s instructions. The E6 mutants pQCXIN-Flag11E6^W133R^, pQCXIN-Flag11E6^L111Q^, pQCXIN-Flag11E6^L70A^, pQCXIN-Flag16E6^L50G^, pQCXIN-Flag16E6^F69A^ were constructed using a KOD-Plus-Mutagenesis Kit (TOYOBO, Japan), prior to DNA sequencing to ensure that no additional base changes were present. The primer sequences used for mutagenesis are listed in supplementary table 1. The E6AP-specific shRNA construct pCL-SI-MSCVpuro-H1R-E6APRi4 was described previously (Handa et al., 2007). pBOB-EF1-FastFUCCI-Puro was a gift from Kevin Brindle & Duncan Jodrell (Addgene plasmid 86849) (Koh et al., 2017).

### Retrovirus and lentivirus transduction

The production and infection of recombinant retroviruses or lentiviruses were accomplished as previously described (Naviaux et al., 1996; Tani et al., 2019). To generate NIKS cells expressing E6, 2×10^5^ cells were seeded in each well of a 6-well plate the day before transduction. Cells were inoculated with viruses at MOI>1 in the presence of 4ug/ml of Polybrene (Santa Cruz). Stable NIKS populations were generated following selection with puromycin (10ug/ml), G418 (50ug/ml) or hygromycin (10ug/ml).

### Organotypic raft culture

Raft cultures were established as previously described (Flores et al., 1999; Lambert et al., 2005). EF-1F human foreskin fibroblasts were mixed at a concentration of 10^7^ cells/ml with Rat Tail Collagen Type I (SLS, 354236) to make the dermal equivalent. Dermal equivalent was allowed to contract in DMEM for four days before NIKS cells were plated on at a density of 1.5 × 10^6^ cells/50ul. Organotypic rafts were firstly cultured in FC media to allow attachment and expansion, followed by cornification media (Flores et al., 1999) to facilitate the formation of cornified layer. Rafts were allowed to stratify for approximately 14 days, then trimmed and fixed in 4% paraformaldehyde (PFA) for 24 hours. Tissue sectioning was performed by histologist at Department of Pathology, Cambridge.

### Immunofluorescence

Immunofluorescence was performed as described previously (Wang et al., 2004). The formalin fixed, paraffin embedded (FFPE) tissue sections were wax removed with Xylene and incubated in Target retrieval solution pH 9.0 (Dako, Glostrup, Denmark) for 10 min at room temperature prior to incubating for 15min at 110°C. The sections or cell samples were washed in PBS and fixed in 4% paraformaldehyde (PFA) in PBS for 10 min at room temperature. Cells were permeabilised in PBS with 0.1% Triton X-100 (Promega) for 30min, then washed in PBS. The sections or cells were blocked in 5% normal goat serum in PBS for 1 hour prior to incubation of the primary antibodies overnight. The antibodies used were listed in supplementary table 1. Antigen antibody complexes were visualised with anti-mouse Alexa 488- or 594-conjugated antibody (Thermo Fisher Scientific) or Immpress anti-mouse/rabbit coupled with tyramide amplification kit (PerkinElmer, Inc). Nuclei were counterstained with DAPI.

### RNA in situ hybridisation

Viral RNA in cells were detected and visualized with RNAscope in situ hybridization assay (Advanced Cell Diagnostics, MN, USA) following the manufacturer’s instructions. The probe used for low-risk E6/E7 RNA detection was RNAscope Probe-HPV6/11 (415211).

### Competition assays

In order to represent the growth condition of the basal layer of stratified epithelium in 2D in vitro assay (Saunders-Wood et al., 2022), NIKS were seeded at high (confluent) density on CellCarrier-96 well ultra Microplates (Perkin Elmer). To each well, 2.4×10^4^ NIKS cells of each mCherry and eGFP were seeded with 6 × 10^3^ irradiated J2-3T3 feeder cells. Cells were cultured for up to 9 days. Media was changed every other day before being fixed in 4% PFA for 30 minutes. Bottom layer and second layer of the cells were visualised and scanned by Harmony Opera Phenix high content imaging system at MRC Institute of Metabolic Science (IMS), Cambridge. Magnification 20x, Field size 0.42mm^2^.

### SDS-PAGE and Western blotting

Proteins were extracted from cells using RIPA buffer and quantified using the BCA protein assay kit (Pierce), before being separated on 4-12% gradient polyacrylamide-SDS-Tris-Tricine denaturing gel (Invitrogen) and transferred onto PVDF membranes (IPFL00010, Merck). After transfer, membranes were blocked for 1 hour at room temperature in 5% milk in TBS. Blots were then incubated overnight at 4 °C with appropriate primary antibody diluted in 5% milk in TBS. This is followed by incubating with appropriate IRDye 800cW fluorescent secondary antibody (Licor) for an hour at room temperature. Protein bands were detected with Odissey imaging system (Licor). Primary antibodies used in this study are listed in supplementary table 1.

### RNA sequencing

Total RNA was extracted from three independent clones of NIKS parental control cells, NIKS-E6AP^-/-^, or NIKS transduced with E6 using the RNeasy mini kit (Qiagen). PolyA selection, reverse transcription, library construction, sequencing and bioinformatics analysis were performed by Novogene. Differentially expressed genes were selected based on a log2(FoldChange) >= 1 & padj <= 0.05 cut-off and were analysed for enriched biological processes using the GO (Gene Ontology) enrichment analysis tool.

### qRT-PCR

Total RNA from NIKS was purified by using an RNeasy Mini Kit (Qiagen), with genomic DNA removed by Turbo DNA-free kit (Invitrogen). cDNA was synthesised with SuperScript III Reverse Transcriptase (Thermo Fisher scientific) using 100uM oligo dT, according to the manufacturer’s instructions. The YAP-responsive genes PTGS2, PLAU, AREG, AXL, TGFBR3 and E6 gene and GAPDH were measured by ViiA 7 Real-Time PCR system (Life Technologies) using Fast SYBR master mix (Applied Biosystems) with 15min denaturation at 95 °C, followed by 45 cycles of 95 °C for 15s and 60 °C for 60s. The PCR primers for qPCR are listed in supplementary table 1.

### Clinical samples and ethics

This study was approved by the institutional review board (IRB, Helsinki Committee) of The Galilee Medical Center. Approval Number NHR 0202-18 on March 12, 2018. The mode of collection, processing, and patient data-handling of the clinical samples used in this study have been described previously (Griffin et al., 2015).

## Acknowledgements

This work is supported by the Medical Research Council (MC-PC-13050 and MR/S024409/1), Chinese scholarship council and Cambridge Trust. We thank Lawrence Banks for his generous gift of px459-E6AP plasmids. We acknowledge Louise Howard for tissue sectioning and the IMS-MRL Imaging Core for high content imaging. We also thank Dr. Yongxu Lu and Qi Zhong for valuable discussions and proofreading the manuscript.

## Author Contributions

Conception and design: WY, NE, JD. Acquisition of data: WY, NE, KZ, AA. Analysis and interpretation of data: WY, NE, HG, JD. Drafting or revising the article: WY, JD.

## Supplementary figures

**Figure 2-supplementary 1.**
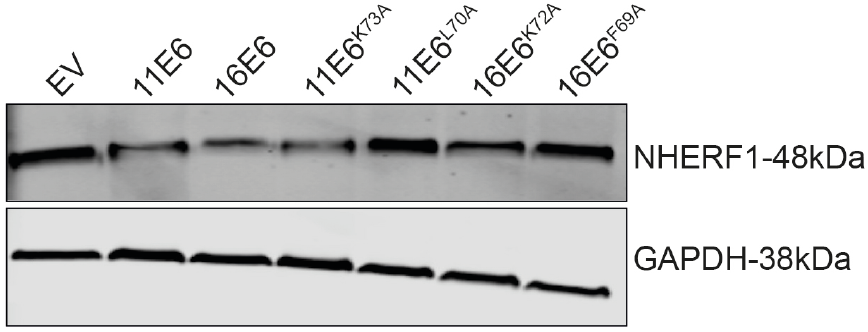
Validation of NHERF1 degradation deficient mutants of E6. NIKS cells were retrovirally transduced with vectors encoding 11E6, 16E6, 11E6^K73A^, 11E6^L70A^, 16E6^K72A^ and 16E6^F69A^. Cells were cultured at 8× 10^6^ cells/well in six-well plates and cell lysates were collected for western blotting. NHERF1 (Santa cruz) and GAPDH (EMD Millipore Corp. USA) primary antibodies were used to detect specific protein bands.

**Figure 4-supplementary 1.**
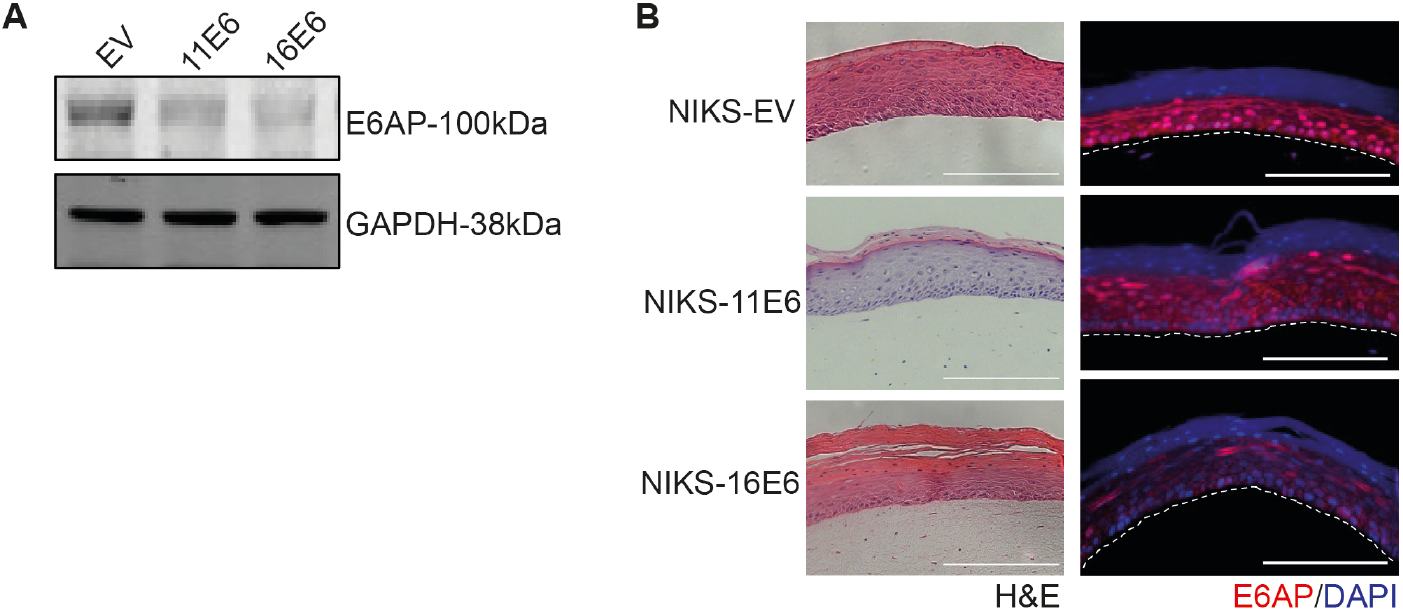
E6 expression causes E6AP level reduction in NIKS. (A) NIKS cells expressing either 11E6 or 16E6 were lysed and subject to western blotting for E6AP (Merck). (B) NIKS cells expressing either 11E6 or 16E6 were used to establish organotypic rafts, followed by immunofluorescent staining with E6AP and DAPI (Merck). Scare bar = 200μm.

**Figure 5-supplementary 1.**
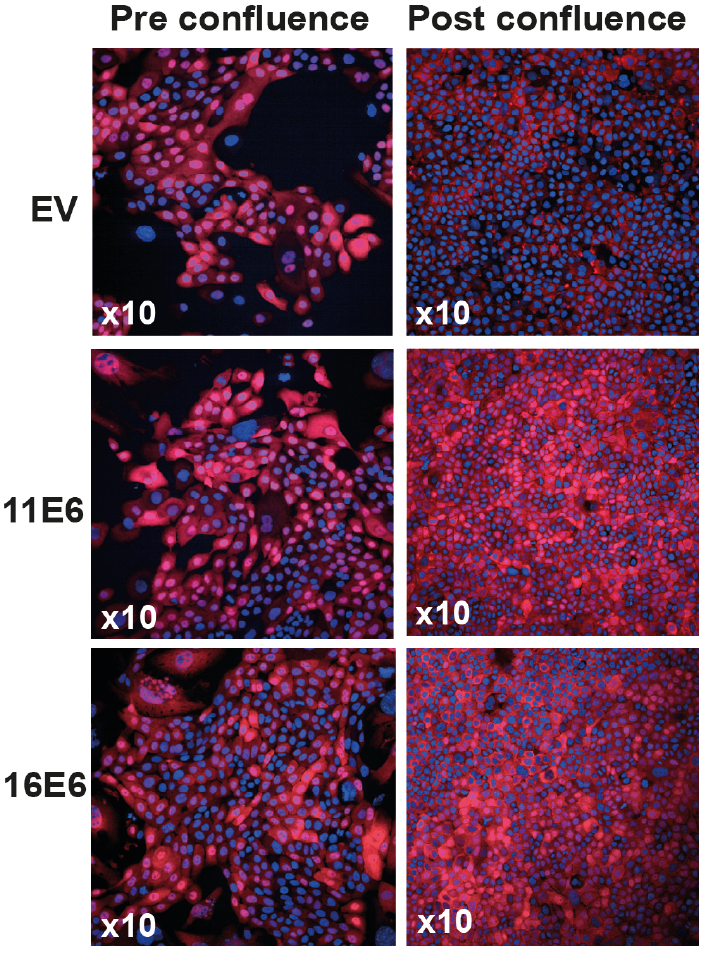
E6 expression promotes YAP nuclear localisation at post-confluence. NIKS cells expressing either 11E6 or 16E6 along with control cells were seeded at 12,000 cells/well (pre-confluence) and 48,000 cells/well (post-confluence) in 96-well plates. Cells were fixed after 72 hours and stained with active YAP antibody (Abcam) and DAPI. Images were captured by Confocal microscope at IMS with 10x magnification.

**Figure 6-supplementary 1.**
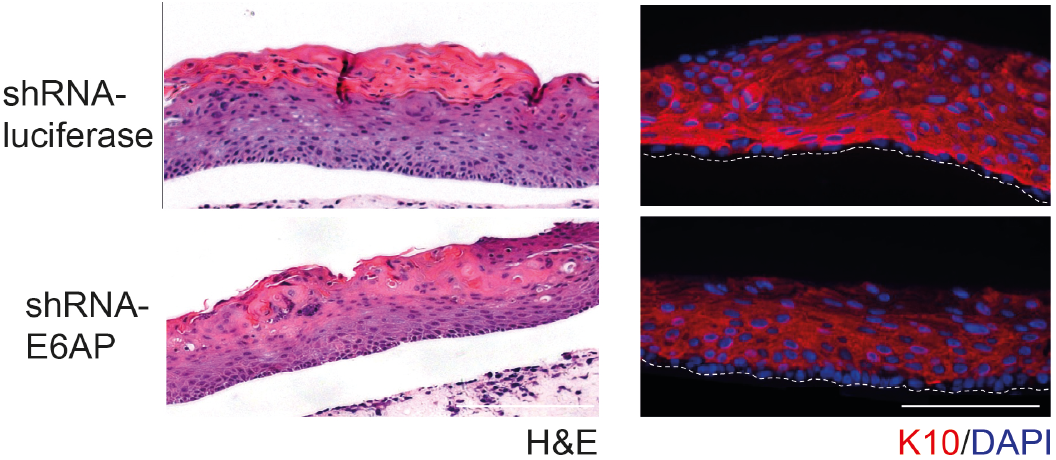
E6AP depletion leads to delay of K10 expression in NIKS rafts. NIKS cells were transduced with lentiviral vectors encoding shRNA targeting either luciferase or E6AP. After selection, NIKS rafts were established, followed by H&E (left) and immunofluorescent staining (right) with K10 (Abcam) and DAPI. Scale bar = 200μm.

**Figure 7-supplementary 1.**
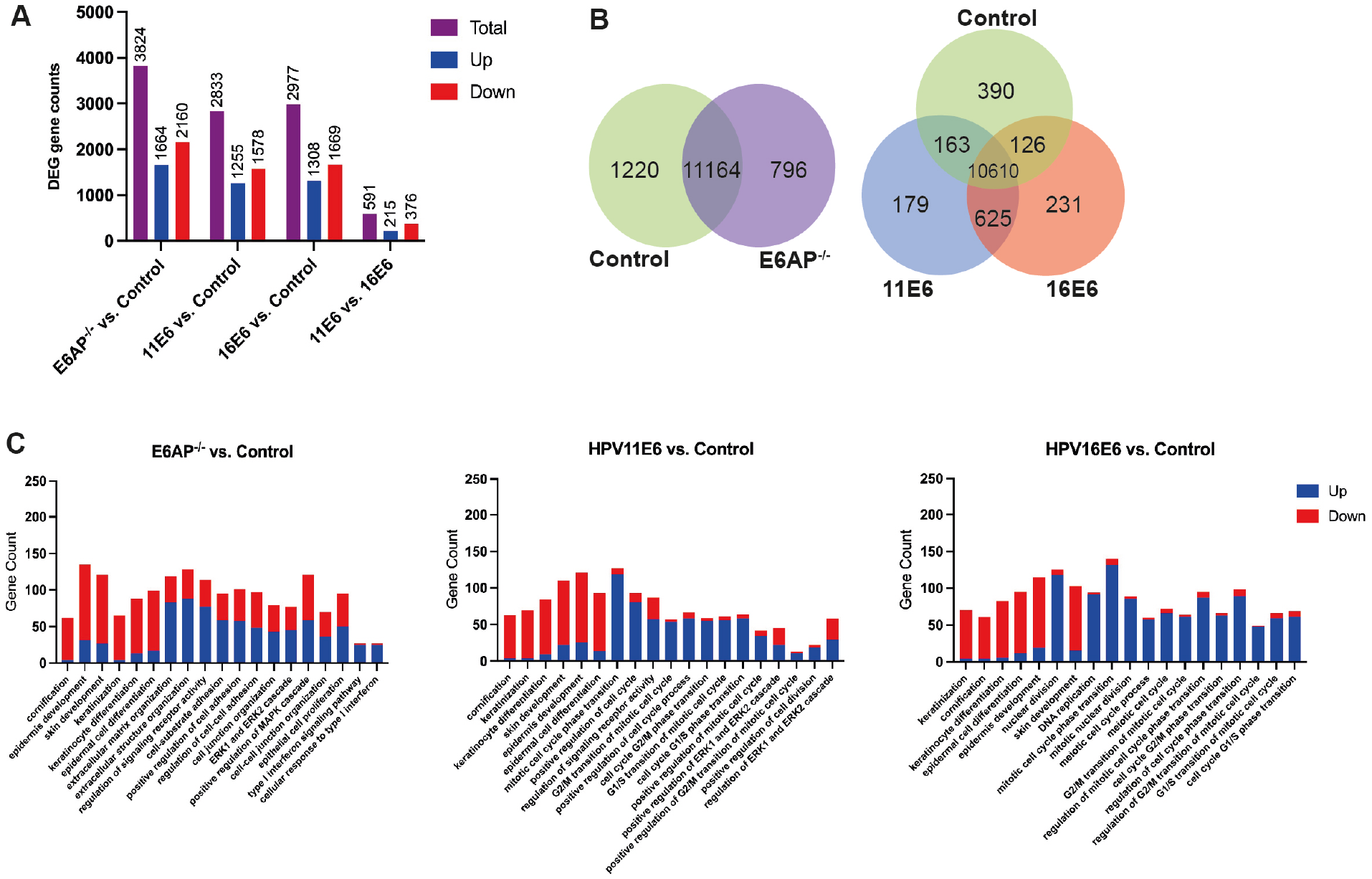
Differential gene expression (DEG) and Gene Ontology (GO) enrichment analysis of RNA sequencing results for E6AP^-/-^ and E6-expressing cells. (A) Total number of differentially expressed genes (DEG) (purple), number of up-regulated genes (blue) and number of down-regulated genes (red) in each experimental group. (B) Vann diagram shows the number of co-expressed genes between the samples. (C) The X-axis displays the selected GO terms that are the most relevant and significant, which ranks from left to right and left is the most significant term. The y-axis shows the number of genes that were up-regulated (blue) or down-regulated (red) under each GO term.

