## Supplementary figures and images for "E6AP is important for HPV E6’s role in regulating epithelial homeostasis and its loss impairs keratinocyte commitment to differentiation"

### heatmap_group_showname E6.pdf

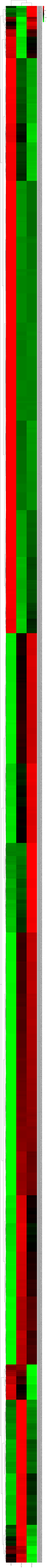

### heatmap_showname E6APKO.pdf

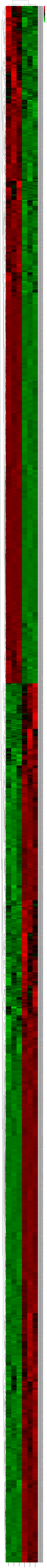
